# Novel inductively-coupled ear-bars (ICEs) for fMRI signal enhancement in rat entorhinal cortex

**DOI:** 10.1101/2022.09.30.510293

**Authors:** Yi Chen, Zachary Fernandez, David C. Zhu, Scott E. Counts, Anne M. Dorrance, Xin Yu, Norman Scheel, Wei Qian, Mahsa Gifani, Chunqi Qian

**Author notes:** **Lead corresponding author:** Dr. Chunqi Qian.

## Abstract

Entorhinal cortex (EC) is a potential target of deep brain stimulation in Alzheimer’s disease (AD) and fMRI can enable whole-brain dynamic mapping noninvasively. However, it remains challenging to study EC-based fMRI connectivity in rodents due to image signal loss and the lower sensitivity of the surface coil ring or array coil for deep brain areas. To reduce the magnetic susceptibility artifacts driven signal loss issue, we introduced baby cream into the middle ear. To improve detection sensitivity, we implemented novel inductively-coupled ear-bars (ICEs) in the 7 T Bruker scanner, which resulted in an approximately 2-fold signal-to-noise ratio (SNR) increase in EC over the conventional surface array. The ICE can be conveniently utilized as an add-on device, with no modulation to the scanner interface. To demonstrate the applicability of ICEs for both task and resting-state (rs) fMRI, whole-brain echo-planar imaging (EPI) was performed in anesthetized rats modeling AD mixed dementia. Seed-based rs-fMRI connectivity maps emanating from the left entorhinal cortex demonstrated its connectivity to the hippocampus, piriform cortex, septal nuclei, and prefrontal cortex. Hence, this work demonstrates an optimized procedure for ICE by acquiring large scale networks emanating from a seed region that was not easily accessible by conventional MRI detectors, enabling better observation of EC-based brain fMRI connectivity studies with a higher signal-to-noise ratio in rodent models of dementia.

## INTRODUCTION

Entorhinal cortex (EC) is a potential target of deep brain stimulation (DBS) in Alzheimer’s disease (AD) patients and animal models of dementia; however, the mechanism for DBS remains unclear. Studies have revealed that DBS of the EC area enhanced memory [1-3] in humans. This is highly consistent with the observation in AD rodent models [4] that provide a platform for more mechanism-based studies by combining optogenetic stimulation with cell-specificity [5-9]. Functional magnetic resonance imaging (fMRI), especially, resting-state fMRI (rs-fMRI) [10], is a useful method to noninvasively study brain-wide dynamics [11] and to provide translational knowledge between humans and animal models. Rs-fMRI has been widely used in both healthy human subjects and patients and across various species with various neurologic, neurosurgical, and psychiatric disorders [12], *e.g*., AD. Despite the promising perspectives to study default-mode network (DMN) abnormalities as a biomarker for AD, EC function/dysfunction within the DMN and its associated network abnormalities are far less explored due to the location of EC in the brain, *i.e*., one of the deepest regions in close proximity to air-containing ear cavities.

The surface coil and coil array could be placed near the rodent brains to have good signal-to-noise ratio (SNR) for the brain parenchyma adjacent to the coil. However, the SNR decreases dramatically as the distance from the coil/coil array increases [13]. This issue is inevitable due to the decaying sensitivity profile of the surface coil/coil array, and especially for rodent imaging inside high-field magnets where the restricted bore is occupied by the rodent holder, air supply tubing, heating pad, etc. To reduce hardware complexity and boost sensitivity, inductively coupled detectors can be placed near the targeted region of interests (ROI) to relay locally detected MR signals with the external surface radiofrequency (RF) coil wirelessly [14-16]. In addition, the sensitivity enhancement in very close proximity to the targeted ROI can be maintained within a distance range, as long as inductive coupling remains larger than circuit loss [16]. Conventionally, teeth and ears of the rodents in the rodent holder were secured by a bite bar and ear bars, respectively [17, 18], in an MRI-compatible stereotaxic frame. The location of EC in the rodent brain [19] makes it practical and feasible to combine the ear bars with inductive coupled detectors, thus enhancing the detection sensitivity.

In addition, magnetic susceptibility artifacts are prevalent on the boundary of air-containing ear cavities in rodents [20, 21]. Although efforts have been made to reduce susceptibility artifacts [22-25], these efforts are mostly focused on restoring lost signals on EPI blood oxygen level-dependent (BOLD) fMRI images over large scales across the entire brain. It still remains challenging to achieve the true whole-brain fMRI imaging. The fMRI image signal loss at the level of EC is due to its proximity to the air-tissue interface in the ears, which introduces a high level of magnetic field inhomogeneity. While the higher magnetic field strength could offer higher signal sensitivity, it also exacerbates the problem of susceptibility artifacts [26]. There is no apparent possibility to use current shims to address the pointillistic character of homogeneity distortion of the magnetic field near the middle ear of rat brain. Several studies have already been done to reduce susceptibility artifact. An equal-TE ultrafast 3D gradient-echo imaging method was developed to provide high tolerance to magnetic susceptibility artifacts in rat cortex with optical fiber implantation at 9.4 T [22]. Since the middle ear filled with air was separated from the external ear by the tympanic membrane [23], this membrane was penetrated and Fomblin Y [23] or toothpaste [24] were used to fill the ear cavities to restore susceptibility-induced MRI signal dropout. However, these methods to diminish the distortion and restore the signal dropout in deep brain regions in proximity to the ear cavities were unsatisfactory because the fluidic Fomblin Y cannot not be kept in the cavities, the toothpaste is skin-irritating, and the implementation procedure needs specific expertise. Therefore, it is highly desirable to have a simple, skin-friendly strategy without modification of MRI interface to restore the signal dropout in the deep brain regions of rodents, including EC, amygdala [27], etc. This will allow a true whole-brain imaging scheme to provide a comprehensive understanding of brain network and function.

Hence, as a proof-of-concept demonstration, we present an animal fMRI strategy for restoring EPI signal loss adjacent to ear cavities and for improving MR detection sensitivity of deep brain regions, in particularly EC, by ICE (Inductively Coupled Ear bars) combined with ear canal injection of baby cream. As an *in vivo* benchmark application of ICE in contrast to conventional surface coil arrays, we have embedded the ICE to show a near 2-fold enhancement of SNR in lateral EC. We evaluated this setup first by using task fMRI with electrical stimulation on the forepaw followed by rs-fMRI. In the task fMRI, we observed a robust evoked BOLD signal along with the ascending projection to the motor cortex from the thalamus. In the rs-fMRI, we observed a robust DMN. Finally, we analyzed the seed-based rs-fMRI connectivity maps based on the left entorhinal cortex (LENT).

## MATERIAL AND METHODS

### Flexible inductively coupled ear bars design and fabrication

Inside an MRI scanner, the rat head is secured by a bite bar from the rostral side and two ear bars from the lateral sides, leaving its dorsal side accessible by surface coils that are more effective for the upper half of the brain. To enhance detection sensitivity of the lateral side in the lower half of the brain, an inductively coupled resonator was integrated with the ear bar to replace the conventional ear bars to stabilize the head during scanning (**Fig.1a**). The ear bar was 3D-printed to have a 9-mm flange and a 2-mm shaft. The shaft was 3-mm away from the center of the flange, so that when the shaft was inserted into the ear cannel and fit inside a positioning hole on the cradle, the center of the flange could be aligned approximately with the ENT region that was about 3-mm above the ear cannel. The flange had a 0.4-mm wide groove around its peripheral edge to accommodate the ICE circuit. As **Fig. 1a** shows, after wrapping a 32-gauge enameled copper wire around the flange for two turns, the conductor leads were self-connected via a zero-biased diode (BBY52, Infenion, Neubiberg, Germany). The junction capacitance of the diode provided necessary capacitance to resonate the circuit around 301 MHz that was slightly above the Larmor frequency of a 7 T scanner, based on the formula: 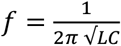. Owing to its proximity with the LENT region, the ICE could sensitively detect regional signals and inductively relay these signals to the surface coil array placed on top of the brain (**Fig. 2d, e**).

**Fig. 1.**
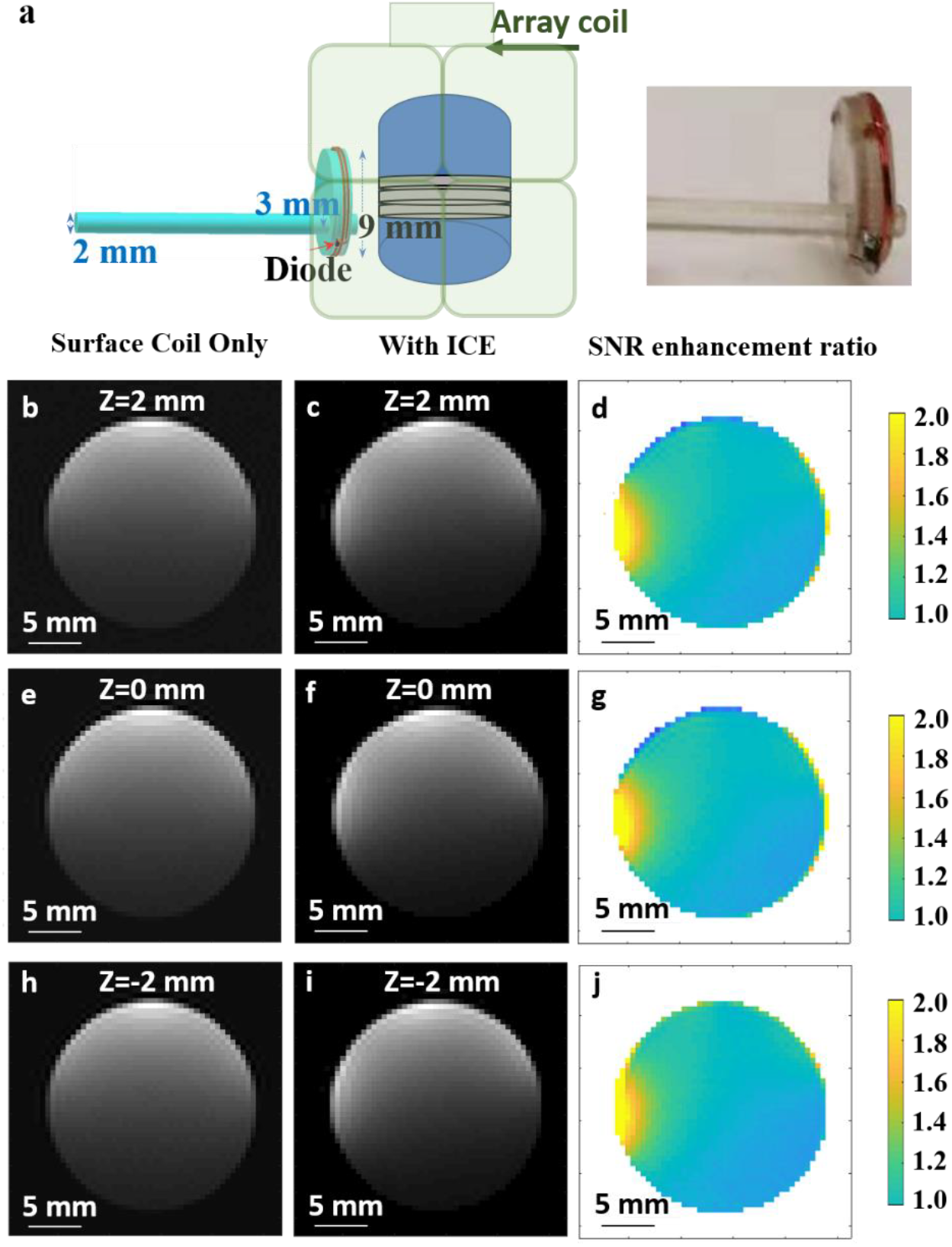
Evaluation of inductive-coupled ear bars with EPI in phantoms. (a) The schematic diagram and the picture of an IDC mounted on the flange of an ear bar for *in vitro* experimental setup. (b-j): The left column is images acquired using the arrayed coil placed above the sample tube. The middle column is images acquired with an additional ICE pressed against the left wall of the sample tube. The right column of SNR enhancement ratio was obtained by dividing images in the middle column with images in the left column. (f) was acquired through the slice passing through the center axis of ICE, while (c) and (i) were acquired through the slices with 2-mm off-sets from the center position.

**Fig. 2.**
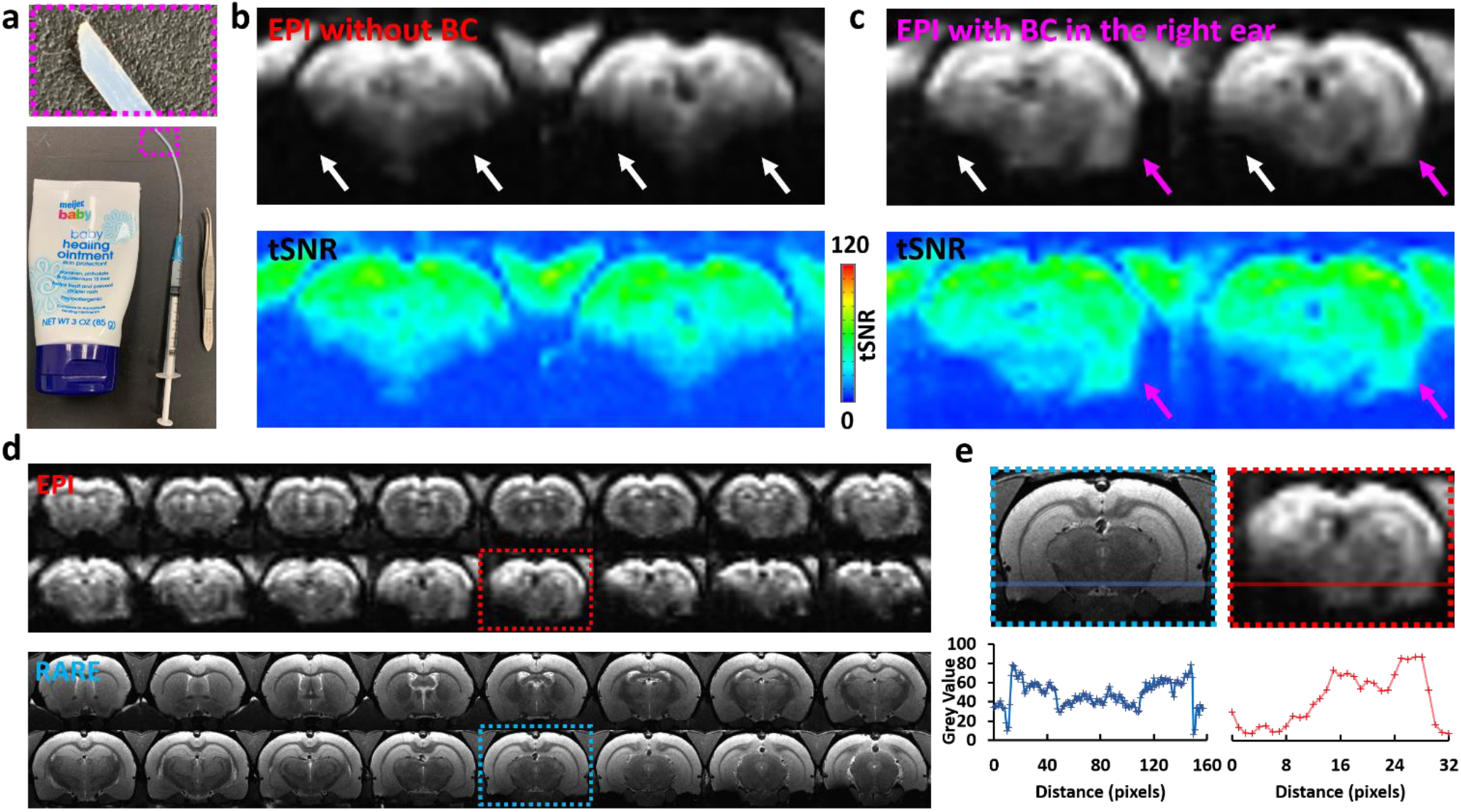
Restored EPI signals in deep brain regions with baby cream. (a) This picture shows the tip of the blunt tubing used to puncture the tympanic membrane and to introduce baby cream into the middle ear. (b) These pictures show representative EPI (upper row) and tSNR maps (lower row) (voxel size = 0.5 × 0.5 × 0.5 mm^3^, TR = 1 s) without baby cream. White arrows demonstrate the susceptibility induced EPI signal loss. (c) These pictures show representative EPI (upper row) and tSNR (lower row) (voxel size = 0.5 × 0.5 × 0.5 mm^3^, TR = 1 s) affected by the baby cream. Magenta arrows demonstrate the restored EPI signals in the right hemisphere with baby cream infusion in the right ear, while white arrows demonstrate the susceptibility induced EPI signal loss in the left hemisphere. (d) These pictures show representative EPI (upper two rows) and RARE images (lower two rows) (pixel size = 0.1 mm × 0.1 mm, thickness 0.5 mm) with baby cream placed in the right ear and none in the left ear. (e) The image signal profile demonstrates the EPI signal restoration in the right hemisphere following baby cream injection, while ear cavity has negligible influence on the image quality of RARE.

### Animals

All procedures in this study were conducted in accordance with guidelines set by the Institutional Animal Care and Use Committee of Michigan State University. We developed a rat model for mixed dementia by breeding the Tg344-19 rat model of AD [28, 29] with the spontaneously hypertensive stroke-prone rat (SHRSP) model of cerebrovascular small vessel disease [30, 31]. Tg344-19 AD rats express the human mutant APPswe and PS1Δ9 genes under control of the mouse prion promoter [28]. Hemizygous Tg344-19 rats were backcrossed onto the SHRSP genetic background and the animals used in this study were 10^th^ generation of progeny. Transgene-positive (Tg+) animals were confirmed by PCR analysis of DNA extracted from ear punches. The present study used a total of 9 Tg+ or Tg-mixed dementia rats (4 male, 5 females) [29, 32]. All animals were three-in-one-housed in 12-12 hour on/off light-dark cycle conditions to assure undisturbed circadian rhythm and ad libitum access to chow and water.

### Animal preparation for fMRI

Animals were first anesthetized with 5% isoflurane in a chamber and baby cream (Meijer, Grand Rapids, MI) was injected into the ear to recover signal loss and reduce noise contamination. Subsequently, the animal was administered by an initial bolus of subcutaneous injection of dexmedetomidine (0.1 mg/kg, NDC 44567-600-04, WG Critical Care, USA). The isoflurane was then discontinued and the animal was transferred to the scanner with dexmedetomidine (0.1 mg/kg/h) delivered subcutaneously. The animal’s body temperature, arterial oxygen saturation level, and respiration rates were monitored and maintained within normal ranges when the animal was inside the scanner. Spontaneous respiration rate typically ranged from 50 to 70 breaths per min during rs-fMRI image acquisition.

### MRI acquisition

All images were acquired with a 7 T/16 cm aperture bore small-animal scanner (Bruker BioSpin, Billerica, MA). A 72-mm quadrature volume coil and a 1H receive-only 2 × 2 brain surface array coil (RF ARR 300 1H R. BR. 2 × 2 RO AD) were used to transmit and receive magnetic resonance signals, respectively.

Functional images (**Fig. 2d, e**) were acquired with a 3D gradient-echo EPI (GE-EPI) sequence with the following parameters: time of echo (TE) = 20 ms, time of repetition (TR) = 1 s, field of view (FOV) = 2.6 cm × 2.6 cm × 1.6 cm, matrix size = 52 × 52 × 32, voxel size = 0.5 mm × 0.5 mm × 0.5 mm. Each rs-fMRI scan acquired 900 time points over 15 mins. We performed electrical stimulation on the left forepaw (5 Hz, 4 s, 333 μs width, 2 mA) (**Fig.3a**) in blocks. Specifically, the block design paradigm included 10-s pre-stimulation, 4-s stimulation, and 11 s intervals, *i.e*., 15 s for each epoch and 8 epochs for a full trial (2 m 10 s), as **Fig. S5a** shows.

**Fig. 3.**
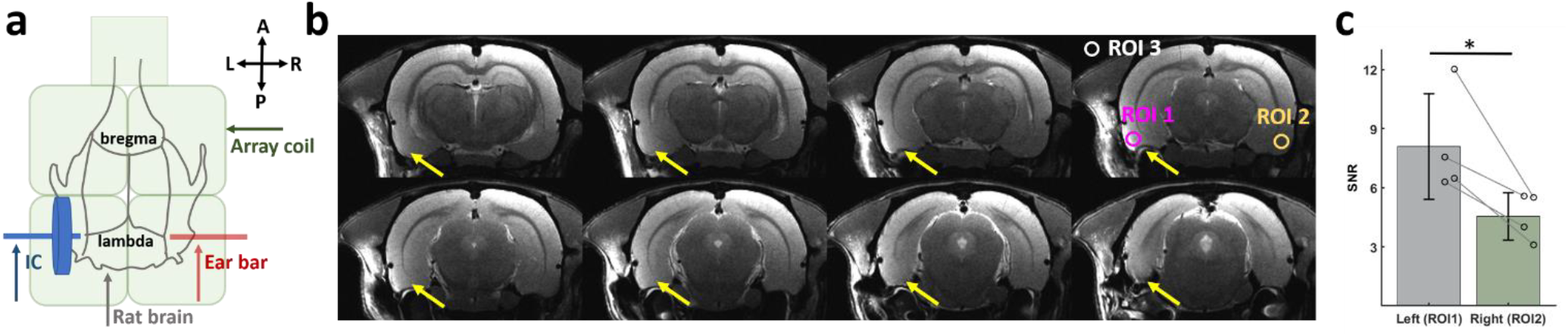
ICE increased the SNR in EC. (a) 2D schematic of ICE placement (blue arrow) in contrast to standard ear bar (red arrow). (b) Example of RARE images with enhanced MRI signal due to ICE in the left hemisphere and without enhancement in the right hemisphere. (c) Significantly increased focal signal intensity in ROI 1 with the ICE placed on the left side compared to the signal intensity in ROI 2 where the right hemisphere has no ICE.

We applied a higher resolution (100 μm) 2D RARE sequence to acquire 32 coronal slices with the same geometry as fMRI images, to accurately identify the LENT in the coronal plane, with the following parameters: TR = 4200 ms, TE = 12 ms, FOV = 2.6 cm × 2.6 cm, matrix = 260 × 260, resolution = 100 μm ×100 μm, slice thickness = 0.5 mm, RARE factor = 4 and signal averaging = 2.

### Immunohistochemistry

Post-fixed brain hemispheres were transferred to 15% sucrose in 0.1 M phosphate buffer until saturated, then 30% sucrose in 0.1 M phosphate buffer until saturated. Brains were frozen on dry ice and sectioned at a 40 μm thickness in 1:12 series in the coronal plane using a freezing-sliding microtome (American Optical, Buffalo, NY). Serial sections were processed for immunohistochemistry using the 6F/3D mouse monoclonal amyloid-beta antibody (1:50; ThermoFisher, Waltham, MA) to visualize amyloid plaque pathology.

### Data analysis

All signal processing and analyses were implemented in MATLAB software (Mathworks, Natick, MA), FMRIB Software Library (FSL) and Analysis of Functional NeuroImages software (AFNI, NIH, USA). For evoked fMRI analysis of **Fig. 2d, e, Fig. S3**, and **S4**, to generate BOLD functional maps, we applied pre-processing steps including motion correction, image registration, time course normalization and averaged fMRI datasets from multiple trials for each animal. The regression analysis of hemodynamic response function (HRF) was based on the BLOCK function of the linear program 3dDeconvolve in AFNI. BLOCK (d, 1) computes a convolution of a square wave of duration d and makes a peak amplitude of block response = 1.

For the resting-state analysis (**Fig. 4d, Fig. 5**, and **Fig. S6**), the flowchart of rs-fMRI processing pipelines for ICA and seed-based analyses is shown in **Fig. S5b**. The preprocessing procedures followed those commonly used protocol in rat rs-fMRI data [33-38], including motion correction, despike, spatial blurring with a full width at half maximum of 0.5 mm, and 0.001-0.4 Hz bandpass filtering in AFNI. Then we conducted ICA analysis with 60 components using MELODIC in FSL to identify and remove non-neural artefacts based on components’ temporal, spatial and spectral features. We manually identified each IC as real signal or noise based on their spatial, temporal, and spectral features. After denoising these components defined as noise, fMRI data from each rat were aligned to the averaged anatomical RARE template. Posterior cingulate cortex and lateral EC were selected as seeds to correlate the whole brain fMRI in **Fig. 4d** and **Fig. 5**, respectively.

**Fig. 4.**
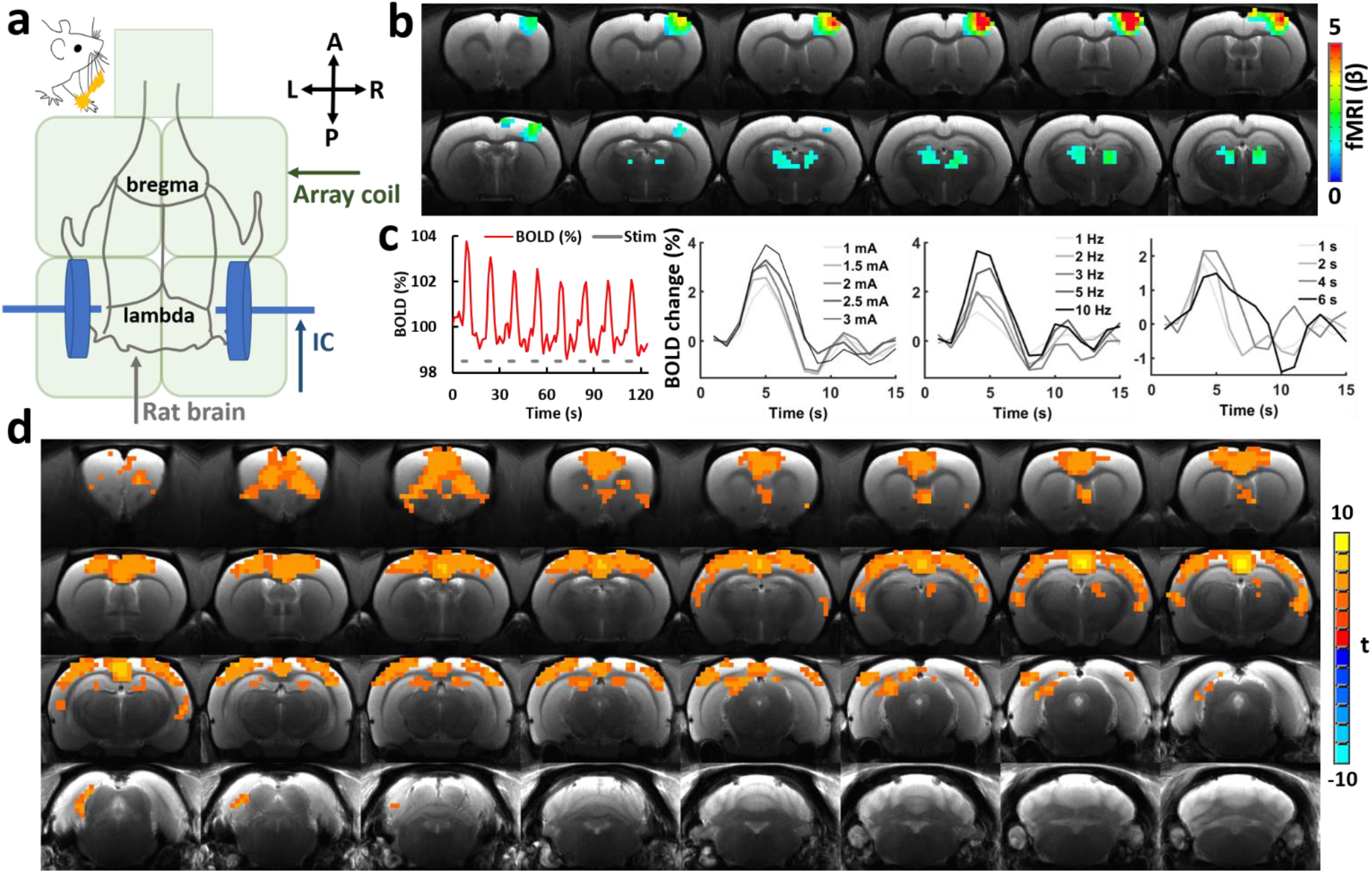
BOLD fMRI responses induced with left forepaw electrical stimulation and DMN rs-fMRI functional connectivity in dementia rats. (a) 2D schematic of bilateral ICE placement (blue arrows) in contrast to standard ear bars. (b) Evoked BOLD fMRI maps of the motor cortex following tactile stimulation of the left forepaw. GLM-based t-statistics in AFNI is used. P (corrected) < 0.005. (c) (left) Averaged tracing of motor cortex BOLD signal evolution over the time course upon forepaw stimulation (n = 5 animals). (right) Mean BOLD signal alterations following various stimulation intensities (1, 1.5, 2, 2.5, 3 mA), frequencies (1, 2, 3, 5, 10 Hz), and durations (1, 2, 4, 6 s). (d). Default-mode network (DMN) shown in color overly on RARE images, as constructed from ICA and seed-based analysis of rs-fMRI data, with ICEs replacing conventional ear bars and with baby cream fulfilled in middle ears.

**Fig. 5.**
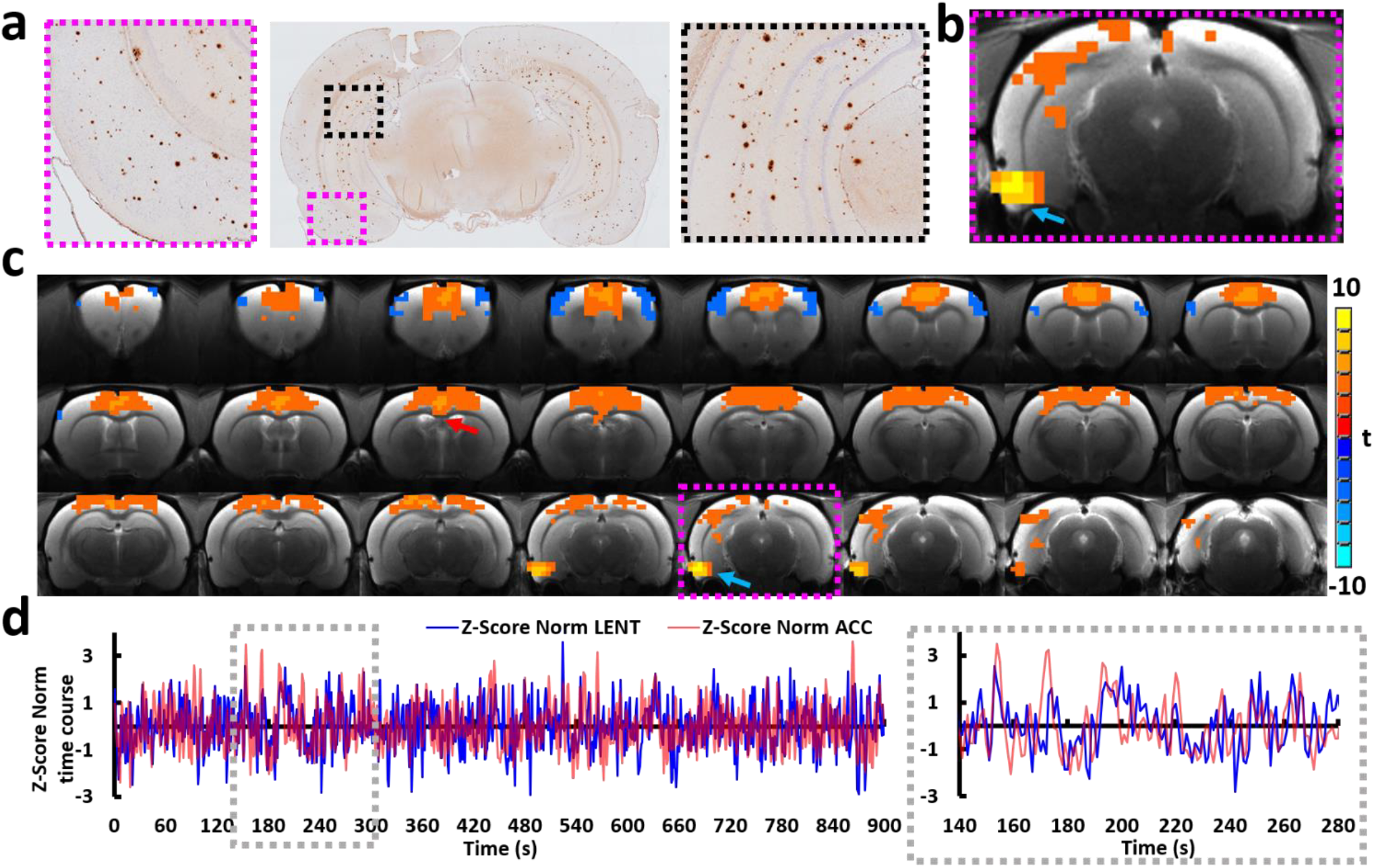
LENT-based connectivity. (a) Amyloid-β deposition with antibodies in the brain slice and anatomical location of the entorhinal cortex in a representative rat. (b) LENT as seed region with ICEs in place, as the blue arrow shows. (c) LENT-based rs-fMRI connectivity maps throughout the whole brain in a mixed dementia rat model (n = 5 animals). (d) Normalized time courses from two highly correlated regions, the LENT (blue line), as the blue arrow shows the ROI in (**c**) and ACC (red line), as the red arrow shows the ROI in (**c**).

**Fig. 2f** illustrates the comparison of SNR by a bar plot containing the enhanced region and the unenhanced region encircled by identical geometries. SNR was calculated by:

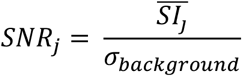

where 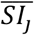 stands for the mean of signal intensity in the regions of interest labeled by *j*, indicating enhanced or unenhanced region, and *σ*_*background*_ stands for the standard deviation of signal intensity in a sample of background. The console of the Bruker system adjusted the RF pulse attenuation automatically for individual rats.

## RESULTS

### Design, characterization, and *ex vivo* evaluation of the ICEs

To emulate the dimension of a rat head, a glass tube with 21-mm diameter was filled by water and secured inside the cradle by ear bars from the contralateral sides (**Fig. 1a**). First, we acquired gradient refocused echo-planar imaging (EPI) of the phantom using phased array coil placed above the sample tube, based on the following parameters: TE = 20 ms, TR = 1 s, FOV = 26 × 26 × 16 mm^3^, Matrix size = 52 × 52 × 32. We repeated the image acquisition twice and calculated the SNR map (**Fig. 1b, c, h**) by dividing the average signal intensity of each voxel with the standard deviation of each voxel in the difference image. Then, we replaced the left ear bar with the ICE and pressed its flange surface against the outer surface of the glass tube. Based on this detection configuration, we acquired another set of images using the same parameters and calculated the SNR map again (**Fig. 1c, f, i**). The signal enhancement capability of the ICE was visualized by the calculating ratio between the SNR maps obtained in the presence and absence of ICE (*i.e*., *Enhancement ratio* = *SNR*_*ICE*_/*SNR*_*No ICE*_). As shown in **Fig. 1d, g, j**, the ICE can enhance detection sensitivity by >2-fold for regions up to 2-mm away from the sample tube’s surface. For a distance separation up to 5-mm from the surface, the ICE can gain at least 20% improvement in detection sensitivity. This effective range is more than enough to cover the EC area in the brain cortex when the ICE is pressed against the rat’s ear. In comparison, the contralateral side of the sample tube was detected only by the surface coil, leading to no sensitivity improvement.

### *In vivo* evaluation of baby cream and ICEs with restored BOLD fMRI mapping in rat brains

After applying baby cream in the ears, ICE was evaluated *in vivo* to measure enhanced MRI signal in the left (**Fig.2**) or right (**Fig.S1**) hemisphere. First, introduction of the baby cream through a blunt tubing puncture into the middle ear allows restoration of GE-EPI signal loss induced magnetic susceptibility mismatch (**Fig. 2a**). Because the baby cream is solid inside the ear, no plug was needed to keep it from leaking out, without additional hardware complexity. **Fig. 2d** shows an example of EPI images (upper row, voxel size = 0.5 mm × 0.5 mm × 0.5 mm, TR = 1 s) and RARE images (lower row, pixel size = 0.1 × 0.1 mm, slice thickness = 0.5 mm). The baby cream was used to fill the right middle ear cavity and ear canal, but none was used in the left ear. Compared to the restored EPI signals in the right hemisphere, it is obvious that part of lateral hypothalamus, the majority of the EC, and almost the entire amygdala in the left hemisphere are affected by the susceptibility artifacts (**Fig.2c**). Noteworthily, both tympanic membranes (eardrum; myringa) were penetrated when the baby cream was injected to the middle cavity and ear canal in both ears, leading to minimum brain activity due to the MRI scanner acoustic noise.

Next, a 9-mm single loop ICE was used to replace the left ear bar beneath the Bruker commercial surface coil array to compare the SNR of deep regions in two hemispheres (**Fig. 3a**). By relaying locally detected MR signals to the external coil array, focal intensity enhancement by this ICE was observed in the left entorhinal cortex from anatomical RARE MR images (**Fig. 3b**, yellow arrows). It is noteworthy that no modifications to the scanner signal interface were required. The focal signal intensity in ROI 1 with the ICE in the left hemisphere was significantly higher than that of ROI 2 in the right hemisphere without ICE (**Fig. 3b, c**), and vice versa when right ear-bar was replaced by the ICE (**Fig. S1a**). It is noteworthy that the ICE could cover a large width along the anterior-posterior direction (particularly for paraflocculus in the cerebellum, **Fig. S1b**) and the ventral-dorsal direction (**Fig. S2**). This setup allows enhanced fMRI signal for restored functional connectivity in rat brains.

### Evaluation of ICE at fMRI platform with electrical stim-driven BOLD mapping and rs-fMRI

Next, after replacing the ear-bars with two ICEs, the setup was evaluated by measuring whole-brain fMRI signal with electrical stimulation on the left forepaw and then rs-fMRI connectivity in anesthetized rats (**Fig. 4**). The brain activation pattern upon left forepaw electrical stimulation is presented in **Fig. 4b**, showing robust BOLD fMRI signal along with the ascending projection to the motor cortex from the thalamus. **Fig. 4c** shows the temporal evolution of the BOLD signals in the motor cortex with the mean time courses acquired at different stimulation intensities, pulse frequencies, and pulse durations. Functional patterns at various stimulation parameters are demonstrated in **Fig. S3,4a**. It is also noteworthy that ICEs ensure highly comparable activation patterns across different animals as shown from five individual rats (**Fig. S3,4a**). **Fig. S4b** shows the activation in the S2, motor cortex in the other hemispheres from two representative rats. For the rs-fMRI, using ICA and seed-based analysis (**Fig. S5**, details in Methods), the DMN is presented in **Fig.4d**, including prelimbic cortex, cingulate cortex, auditory cortex, posterior parietal cortex, and retrosplenial cortex. These results provide strong evidence for highly reliable detection of BOLD fMRI signals under experimental conditions, which are highly consistent with reports from other researchers [37-42].

### Seed-based correlation map of the left entorhinal cortex with ICE for AD animal studies

To demonstrate the advantage of novel ICEs for animal studies, we analyzed the seed-based rs-fMRI connectivity maps of the left entorhinal cortex (LENT) in rats. As shown in **Fig.5a, b**, the EC is located at the caudal end of the temporal lobe in rodents [19]. Due to its diverse heterogeneous projection, we first confirmed amyloid-β deposition with antibodies in the rat brain slices in this study. **Fig. 5a** shows a representative coronal section. Then we chose the region with prominent amyloid-β deposition as the seed to correlate the voxel-wise fMRI signals in a whole brain scale. Highly consistent with other studies [43, 44], we observed correlations in the hippocampus, piriform cortex, septal nuclei, and prefrontal cortex (**Fig. 5c**). **Fig. 5d** shows the Z-score normalized correlations that are highly correlated from the LENT to the anterior cingulate cortex (ACC). It is noteworthy that the correlation maps are different when we choose different subregions of the LENT as seeds (**Fig. S6**), this may result from the heterogeneous projections of EC. These different seed-based whole-brain connectivity patterns from the LENT in rats warrant further studies on the dysfunction of the EC in AD.

## DISCUSSION

Extensive studies have revealed the critical role that EC plays in AD, while functional characterization of EC has been limited by the lack of effective imaging modalities for *in vivo* whole-brain imaging with high SNR in deep brain regions. From the molecular perspective, Khan U. et al have found that the LENT was particularly sensitive to tau and amyloid pathology, and that LENT dysfunction could spread to the parietal cortex [19]. From a structural perspective, Jessen F. et al reported a mean EC volume reduction of 18% in patients with subjective memory impairment, 26% in patients with mild cognitive impairment, and 44% in the patients with AD [45]. Based on a layer specific study, Kobro-Flatmoen A. et al concluded recently that EC layer II contains reelin-positive projection neurons, while there is a linkage between reelin with plaque and tangle pathologies [46]. However, due to the larger distance separating deep brain regions and the surface coil placed above the skull, as well as susceptibility artifacts induced by the air in the ear cavity, it remains extremely challenging to study the functional role the EC plays in AD using fMRI. The EC is less explored in AD animals compared to the widely investigated DMN [47-50], cortex, or subcortical regions, *e.g*., hippocampus. The imaging scheme of ICE with baby cream makes it possible to establish EC-based connectivity in animals (**Fig. 5**), bringing a missing piece to study the EC-mediated whole brain connectivity in AD rodent models. Our methodology may provide new insights into different EC connectivity patterns in healthy and AD animals, enabling us to search for novel preclinical fMRI biomarkers for AD.

We developed the ICE as a way to increase fMRI signal sensitivity. Our strategy has several advantages for rodent brain research. First, it is noteworthy that the baby cream allows easier access and matching solidity to effectively reduce susceptibility artifacts. This strategy provides a true whole-brain mapping platform to noninvasively decipher the functional connectivity dynamics using fMRI, including deep brain regions such as EC and amygdala [51]. Second, the BOLD fMRI with lower structure contrast and spatial resolution are registered to the high-resolution anatomical images for group analysis. With the restored signal dropout, the whole-brain boundary could be used as a landmark to co-register activation maps [52]. This is particularly helpful to keep signals after averaging in the small nuclei that are affected dramatically by the quality of image registration. Without this highly desirable feature, image registration becomes less accurate. Third, as demonstrated in our previous study, surface coils can be combined with inductively-coupled detectors (ICD) and therefore can accommodate for optogenetics with optical fibers passing through the open area of the surface coil and ICD [16], which will enable optogenetic modulation of EC with cellular and circuit specificity in rodents, *e.g*., interneurons, astrocytes etc. To decipher the contribution of specific cell types to destabilizing effects within neural networks and to study the mechanism of the EC as the potential target for deep brain stimulation to treat AD [4, 53], the ICDs show a sensitivity advantage over conventional MRI coils for optogenetic-driven functional mapping. Finally, as shown in **Fig. 3b** and **Fig. S2**, the ICE could increase the SNR across a range of brain regions, and notably, without any modification of MRI scanner interface. When the ICE is applied to various transgenic AD rodent lines [54-58] for awake animal fMRI studies [59-63], it will provide a highly promising imaging platform for awake fMRI in both healthy and diseased animal models.

Several limitations about the usage of ICE should be considered when interpreting the results of this study and for future optimization of the ICE for animal fMRI. First, based on our empirical evidence, if the rat’s body weight is larger than 400 g, which means a larger brain size and larger distance between the brain parenchyma and the ICE, the ICE with a bigger diameter should be considered since the signal enhancement is achieved within a limited ROI when the ICE has a diameter of 9 mm (**Fig.1**). For mouse studies, it will be necessary to re-design a different ICE with matching geometrical parameters. Second, because the materials used for the 3D printing is UV curable photopolymer, it could not be as stiff as the ear bars made of carbon fiber. Therefore, for awake animal studies, there is still room for improvement from the material perspective. It is also worth pointing out that in rare occasions if a rat wakes up during scanning and struggles violently, the ICE can be bent and broken. Thus, measurements were performed only under a well-working anesthetic condition. Third, once the tympanic membranes were penetrated, the baby cream in the ear cavity is difficult to completely remove after experiments. The residual cream could attenuate non-BOLD effect induced by EPI gradient switching sounds. Therefore, this effect should be taken into consideration for longitudinal studies. Last, although the ICE has a 2-fold and obvious SNR enhancement in regions adjacent to the skull, such as the EC and the paraflocculus in the cerebellum, its passive coupling limits the effective enhancement depth, leading to no SNR improvement in regions farther away from the skull, such as the lateral hypothalamus, ventral tegmental area, etc. In the future, it is worthwhile to explore the Wirelessly Amplified NMR Detector (WAND) with the ear bar to actively amplify MRI signals, leading to improved effective detection range with an additional >3-fold sensitivity gain over the passive coupling [64, 65].

## CONCLUSIONS

In summary, we have implemented novel ICEs in the 7 T scanner to yield high-resolution structural and functional images of the rat brain without susceptibility induced signal loss in functional MRI. The ICE-based mapping scheme enables EC-driven brain connectivity studies with a simplified experimental setup, providing opportunities to further study EC in AD animal models using fMRI coupled with additional modilities such as optogenetics and calcium recordings.

## Data availability

The data that support the findings of this study are available from the corresponding authors upon request.

## Code availability

The related image processing codes are available from the corresponding authors upon request.

## ACKNOWLEDGEMENTS

This research was supported by NIH RF1NS113278-01, RF1NS128611-01 (XY and CQ), NIH R01AG060731-A1 (SEC), NIH R21AG074514-01(AMD), NIH R01-HL-13769401 (AMD), R01AG057571 (DCZ) and by the Division of Electrical, Communications and Cyber Systems of the National Science Foundation under award number 2144138 (CQ). This project has also received funding from the European Union Framework Program for Research and Innovation Horizon 2020 (2014-2020) under the Marie Skłodowska-Curie Grant Agreement No.896245. Any opinions, findings, and conclusions or recommendations expressed in this material are those of the author(s) and do not necessarily reflect the views of the funding agencies.

## Supplementary Information

**Figure S1.**
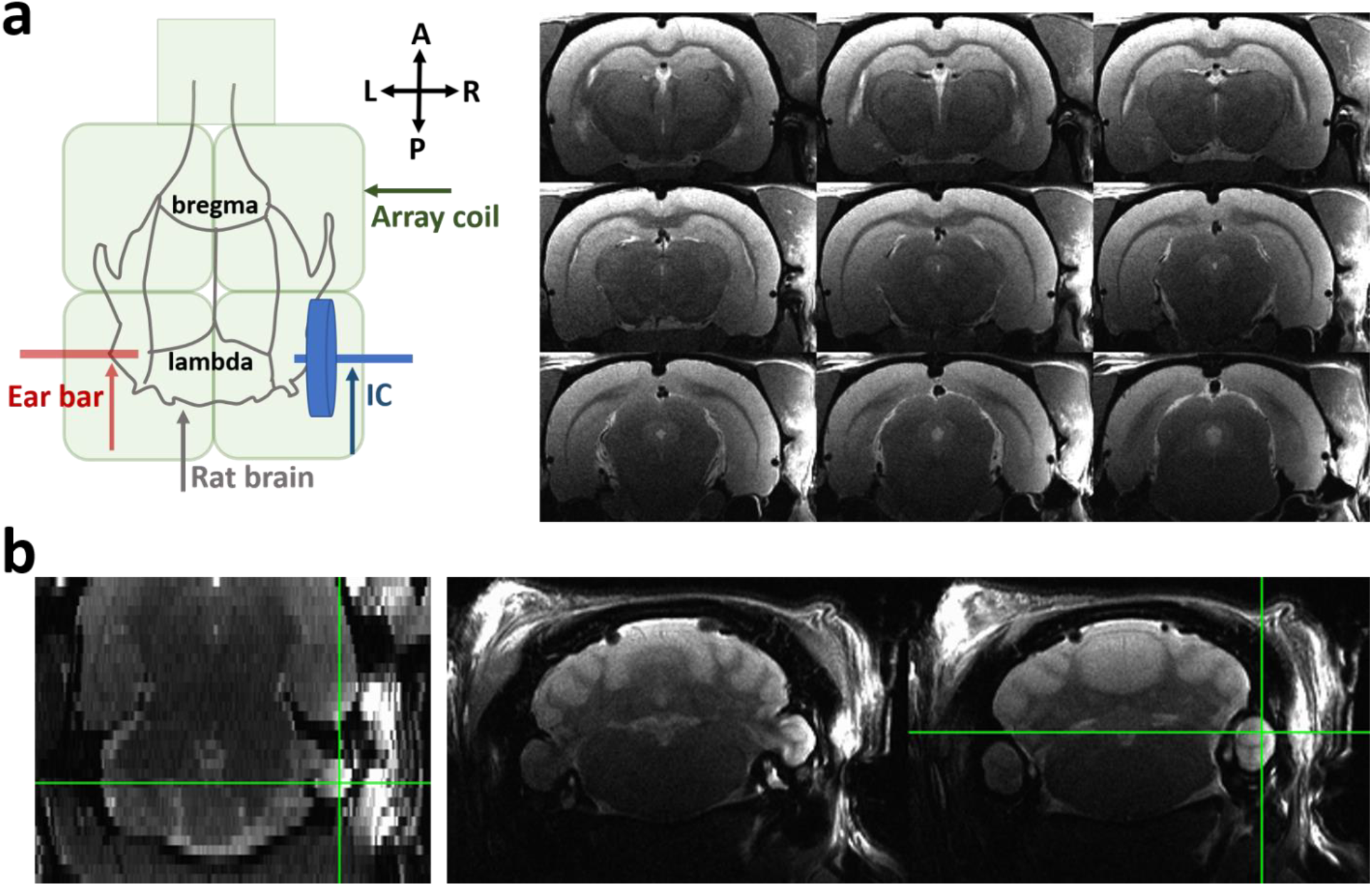
Overview of the inductively coupled ear-bars (ICEs) placement and its application in neuroimaging studies. (**a)** (left) 2D schematic of ICE placement (blue arrow) in contrast to standard ear bar (red arrow). (right) Exampe of an EPI image (voxel size = 0.5 mm × 0.5 mm × 0.5 mm, TR = 1 s) with enhanced image signal due to ICE in the right hemiphere and none in the left. (**b)** ICE could cover a large ROI from the anterior-posterior direction, in particular for paraflocculus in the cerebellum.

**Figure S2.**
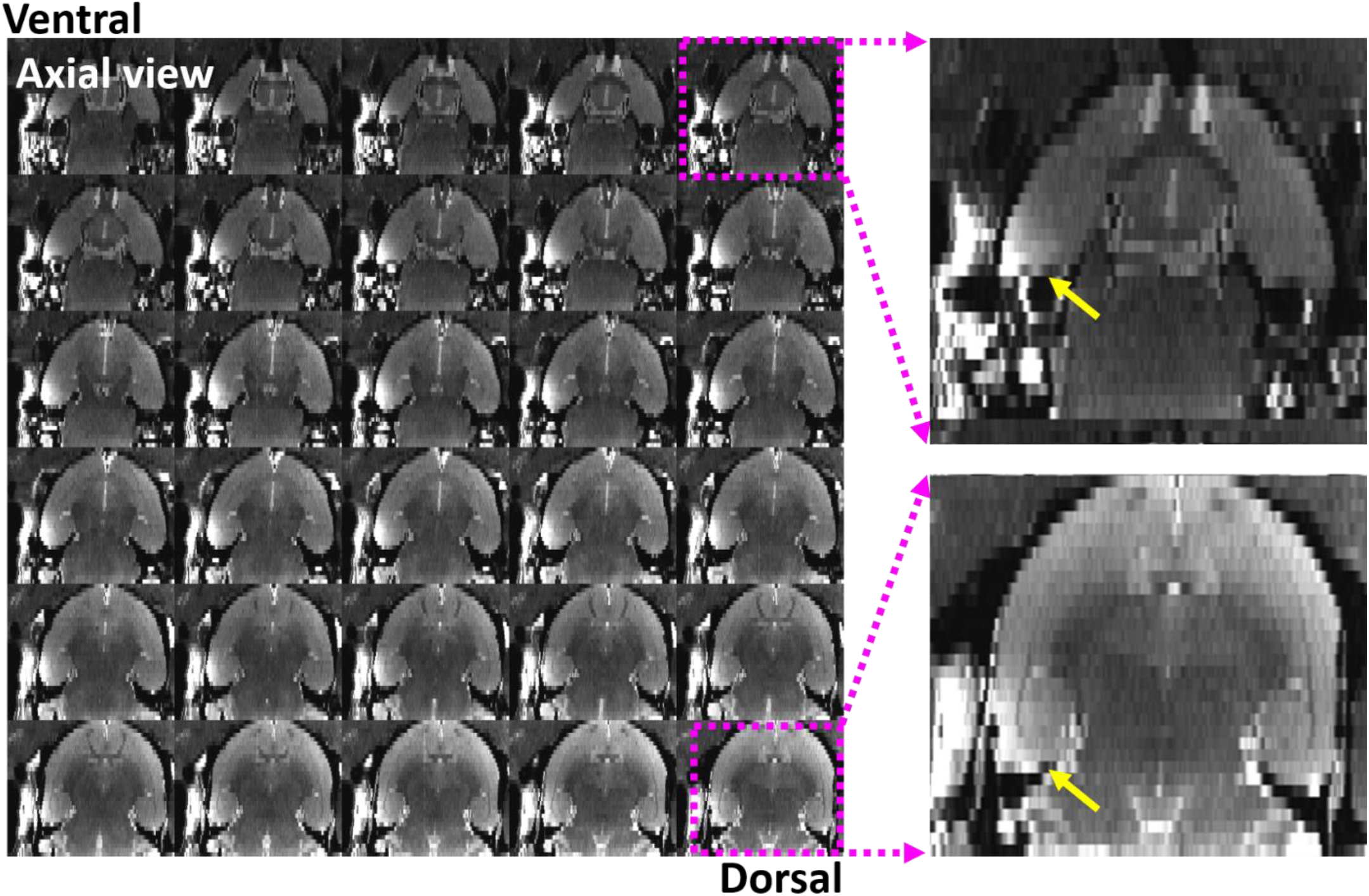
Example image in the ventral-dorsal direction showing ICE is capable of covering large ROIs in this aspect, especially for entorhinal cortex, therefore allowing signal enhancement for restored EPI-based functional signals to evaluate functional connectivity.

**Figure S3.**
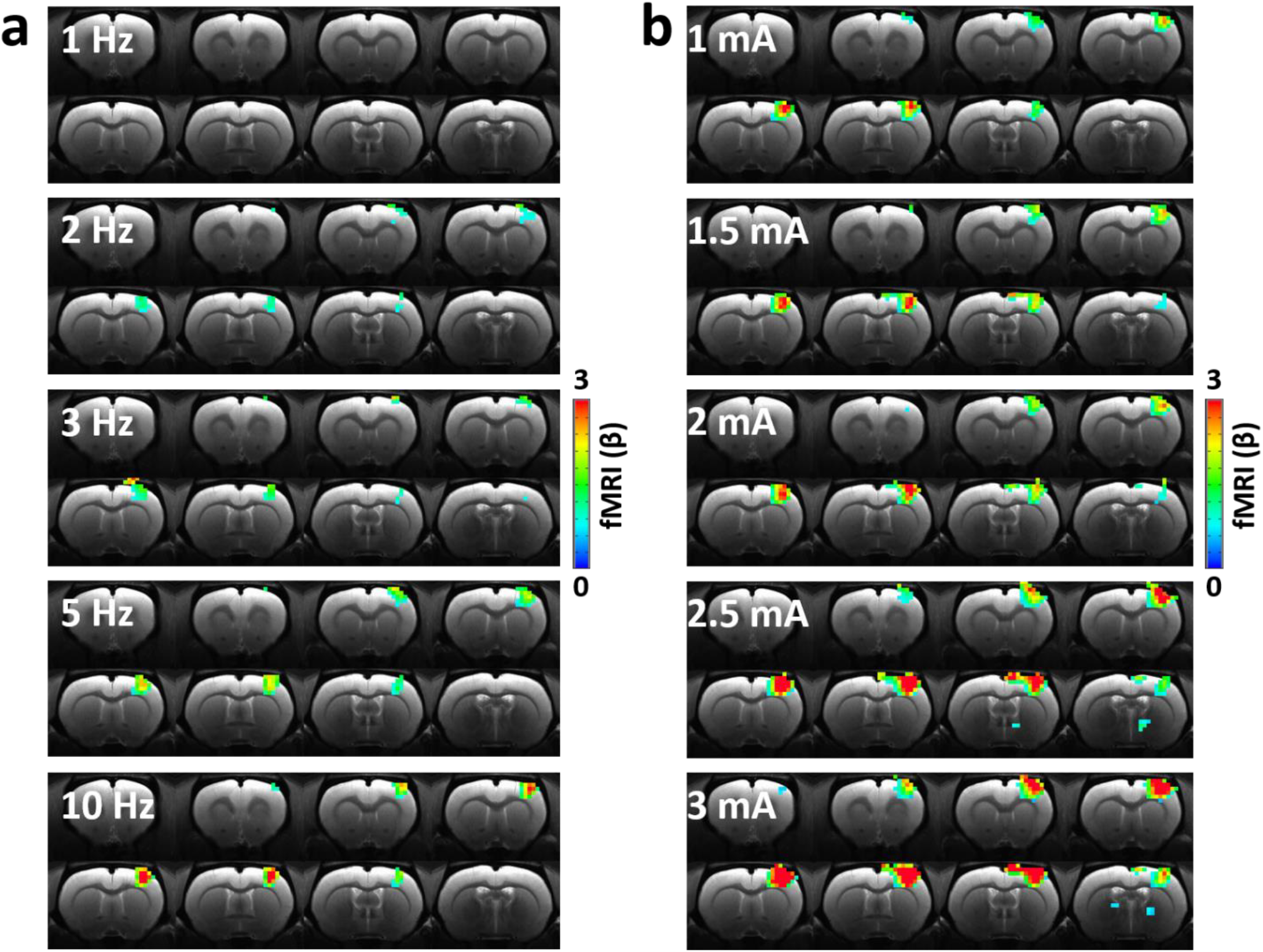
With ICEs in place, shown are evoked BOLD fMRI maps of the motor cortex following tactile stimulation (**a)** with frequencies increased from 1 to 10 Hz, and (**b)** with intensities increased from 1 to 3 mA.GLM-based t-statistics in AFNI is used. P (corrected) < 0.005.

**Figure S4.**
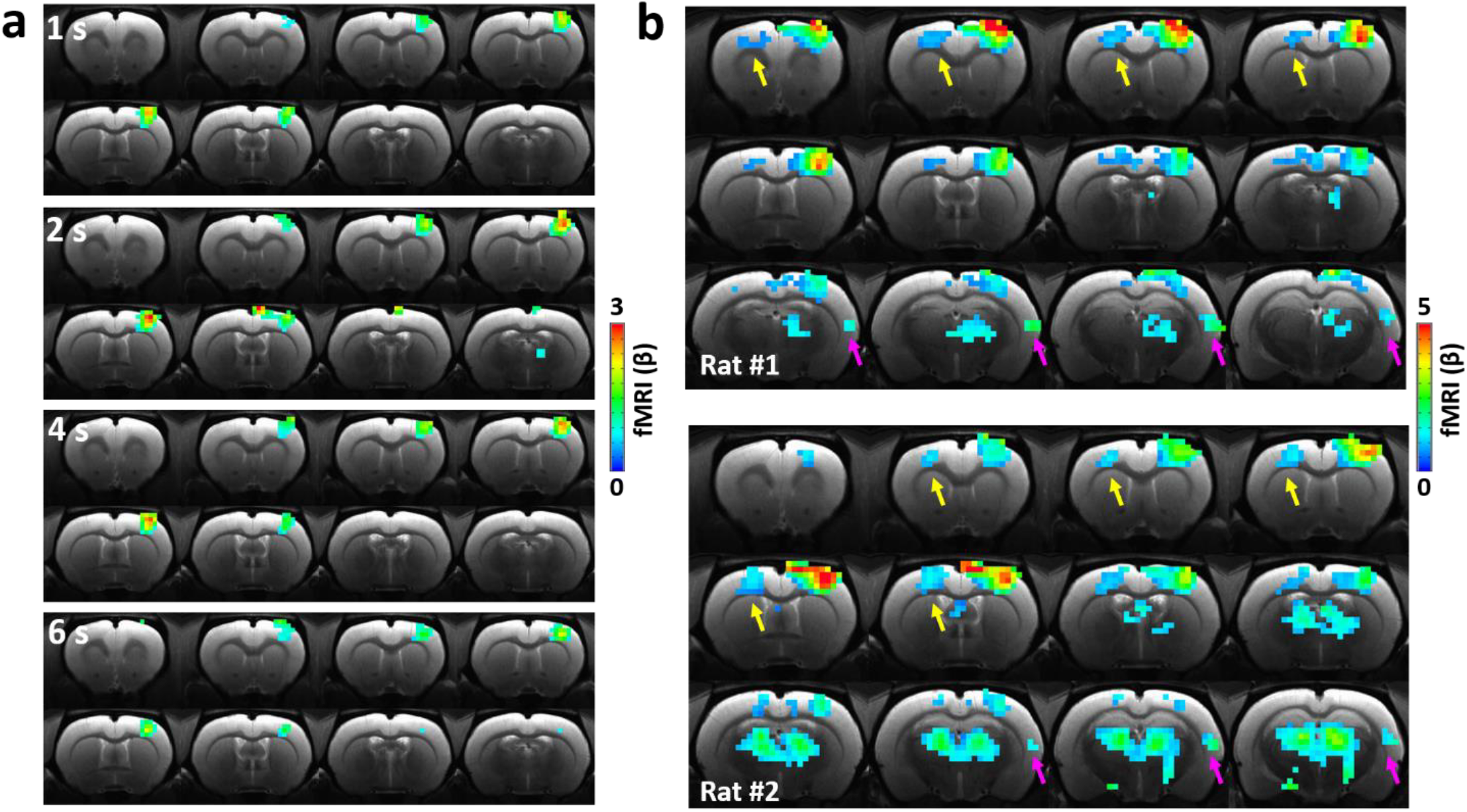
(**a**) With ICEs in place, shown are evoked BOLD fMRI maps of the motor cortex following tactile stimulation with durations from 1 to 6 seconds. GLM-based t-statistics in AFNI is used. P (corrected) < 0.005. (**b**) With ICE, well defined activation maps are shown in the secondary somatosensory cortex (S2, magenta arrows) and interhemispheric motor cortex (yellow arrows) in two representative rats. GLM-based t-statistics in AFNI is used. P (corrected) < 0.005.

**Figure S5.**
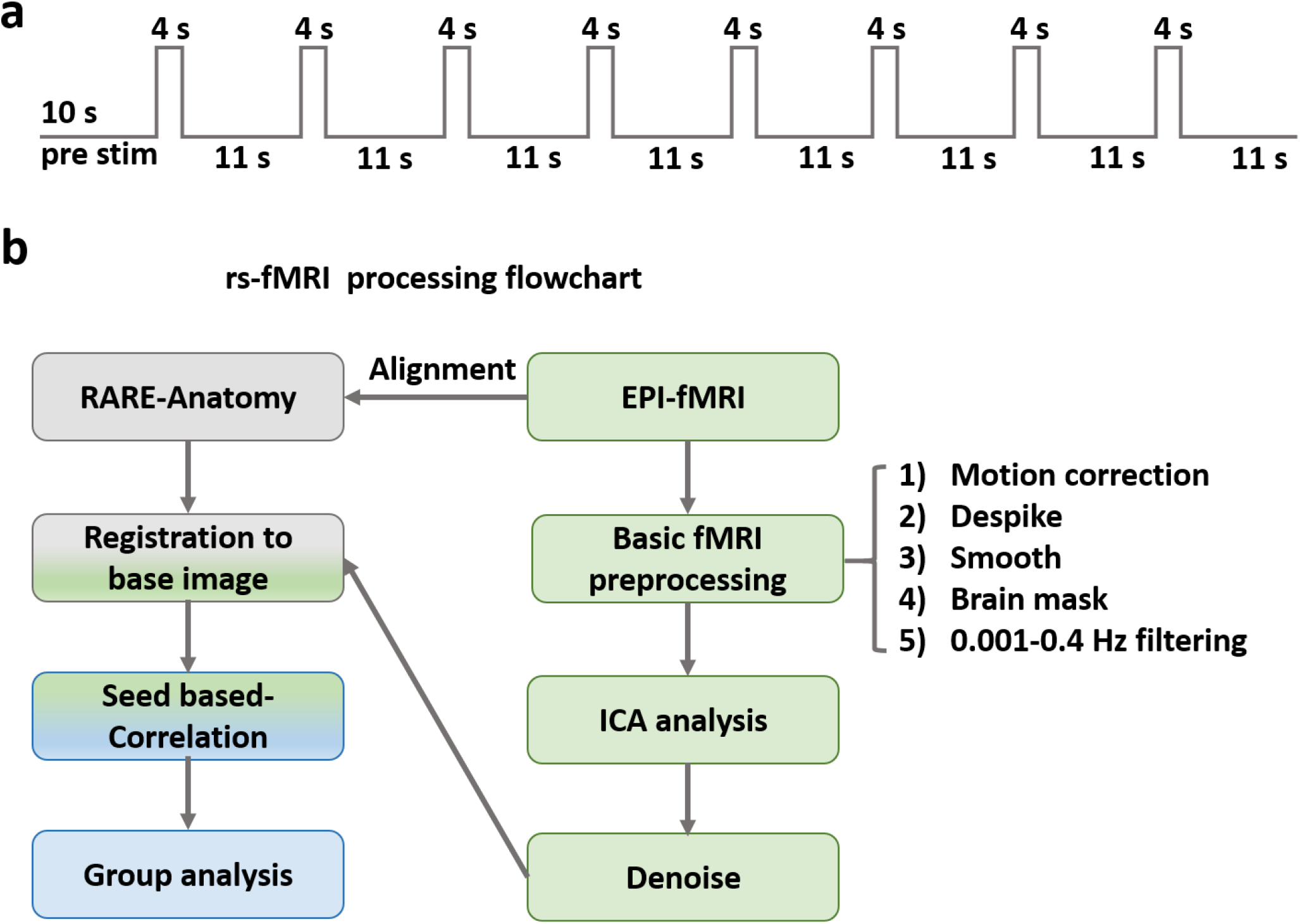
(**a**) BLOCK design paradigm for the task fMRI. (**b**) Flowchart of rs-fMRI processing pipeline for ICA and seed-based analyses.

**Figure S6.**
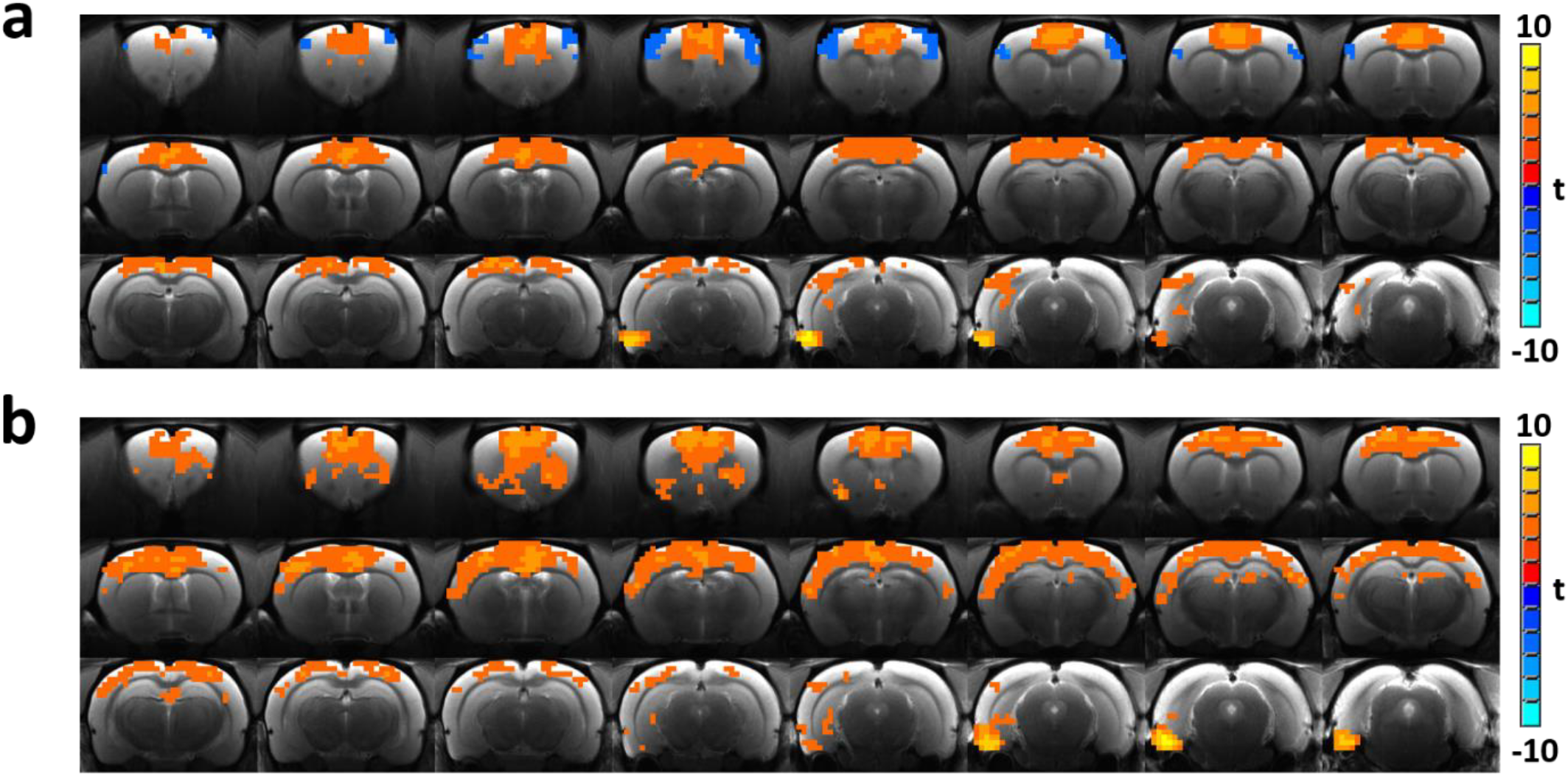
ICE allows us to distinguish different ENT-based functional connectivity correlation maps depending on which subregion is used to place the seed (p < 0.001).

